# Evaluating an internet-delivered fear conditioning and extinction protocol using response times and affective ratings

**DOI:** 10.1101/2021.09.15.460070

**Authors:** Johannes Björkstrand, Daniel S. Pine, Andreas Frick

## Abstract

Pavlovian fear conditioning is widely used to study mechanisms of fear learning, but high-throughput studies are hampered by the labor-intensive nature of examining participants in the lab. To circumvent this bottle-neck, fear conditioning tasks have been developed for remote delivery. Previous studies have examined remotely delivered fear conditioning protocols using expectancy and affective ratings. Here we replicate and extend these findings using an internet-delivered version of the Screaming Lady paradigm, evaluating the effects on negative affective ratings and response time to an auditory probe during stimulus presentations. In a sample of 80 adults, we observed clear evidence of both fear acquisition and extinction using affective ratings. Response times were faster when probed early, but not later, during presentation of stimuli paired with an aversive scream. The response time findings are at odds with previous lab-based studies showing slower responses to threat-predicting cues. The findings underscore the feasibility of employing remotely delivered fear conditioning paradigms with affective ratings as outcome, and highlight the need for further research examining optimal parameters for concurrent response time measures or alternate modes of non-verbal estimation of conditioned responses in Pavlovian conditioning protocols.

## Introduction

Fear conditioning and extinction are widely used experimental protocols in behavioral sciences important for studying associative learning and anxiety ^1,2^. Processes related to threat-induced associative learning is generally held to be theoretically important for understanding the emergence and treatment of anxiety- and trauma-related psychopathology^3^. The popularity of the fear conditioning paradigm is partly due to the possibility to translate findings between humans and animals, which has provided significant progress into the underlying neurobiological mechanisms of associative fear learning ^4,5^.

During fear conditioning, a neutral cue (for example sounds, geometrical figures or pictures of human faces) is paired with an aversive stimulus (US), often an electrical shock or an aversive sound ^2^. Through associative learning, the previously neutral cue becomes a conditioned stimulus (CS+) predictive of the aversive outcome, and starts to elicit conditioned defensive responses (CRs). This is adaptive in an evolutionary sense, but exaggerated threat reactions can contribute to pathological anxiety ^6,7^. In human research, experimental protocols typically include a control stimulus (CS−) never paired with the aversive outcome and often conceptualized as a conditioned safety signal ^2^. Initially considered a neutral control, response to the CS- is of theoretical interest as a measure of safety learning or conditioned inhibition ^8^. During extinction, the CSs are presented again, but omitting the US, typically leading to a decrease in CR expression. Of relevance for clinical anxiety disorders, deficits in extinction have been related to anxiety-related psychopathology ^6^, and trait-anxiety ^9^. Furthermore, extinction is commonly used as an experimental model for exposure-based psychological treatment and lab-based extinction studies can be translated to novel treatment interventions in clinical samples ^10,11^.

To study fear acquisition and extinction, the CR needs to be quantified, and for this purpose various methods have been used. In lab-based studies, the CR is most often quantified through freezing behavior in rodents, and in humans through psychophysiological measurements, such as skin conductance responses (SCR), fear-potentiated startle (FPS), hear-rate changes or pupil dilation, and less commonly through response times. Additionally, self-report based measures are often used, most commonly online ratings of US-expectancy or affective ratings of experimental stimuli pre and post learning phases ^2,12,13^.

Although lab-based studies can further our understanding of fear and extinction learning, they are both costly and work intensive, requiring on-site testing with specialized lab equipment, which is a major impediment for studies requiring large sample sizes. This problem could be mitigated by developing remotely-delivered experimental protocols. Emerging evidence supports the feasibility of such an approach. Through a dedicated smart-phone app using an aversive sound as US and self-report based outcome measures to quantify learning, i.e. US-expectancy and affective ratings, Purves et al. ^14^ were able to demonstrate both fear conditioning, extinction and generalization, with results comparable to in-lab testing. Furthermore, McGregor et al ^15^, used the same methodology in a large sample demonstrating an association between anxiety-related psychopathology and US-expectancy rating during fear and extinction learning. These results are promising, but there is a need for further studies in this area. Firstly, Purves et al.^14^ only included affective ratings prior to acquisition and after the extinction phase. Although the results demonstrated conditioning, they could not demonstrate an effect of extinction using this outcome measure and experimental design. Moreover, although this study found an association between affective ratings and indices of trait anxiety ^14^, the design precluded examinations of independent contributions of acquisition and extinction to anxiety ^6^. Secondly, the protocol developed by Purves et al. uses only self-report-based indices as outcome measures. Although verbal and non-verbal indices of fear learning are correlated, these association are weak, and the measures likely reflect different aspects of the fear response ^12^. Including both would therefore allow for studies capturing broader aspects of the human fear response. Developing non-verbal indices of remotely-delivered fear and extinction learning could arguably reduce the risk of results being influenced by effects related to response bias and demand characteristics, and could possibly be a way to study non-conscious aspects of these processes. Although psychophysiological measurements require specialized equipment, several previous studies have shown that response times can be used as a learning index in fear and extinction studies, showing effects of both acquisition, extinction and reinstatement ^13^. Specifically, previous lab-based studies using concurrent measures of reaction time have shown slower response times to an auditory probe during CS+ presentation compared to CS− ^16,17^. This has been interpreted as an effect of attention capture by the CS+ and could thus be used as a learning index for fear conditioning and extinction. Since existing internet-based research tools that allow for reliable assessment of response time are freely available ^18–20^, this is a promising avenue for developing non-verbal based fear learning indices suitable for remote delivery.

Hence, the purpose of this study was to replicate and extend previous results of remotely delivered fear and extinction protocols, firstly by including measures of affective ratings of CS both after acquisition, extinction, and a reinstatement-test, as well as adding a follow-up test approximately 7 weeks after the original experiment to asses long term retention of this type of learning, and possible effects of spontaneous recovery. Secondly, by adding concurrent measures of reaction time during CS presentation, we aimed to assess a non-verbal index of fear and extinction learning.

## Methods

### Participants

Eighty adults (61 women, 14 men, 5 other/do not want to specify; mean (SD) age: 35.5 (13.5), range: 18-71 years) were recruited from the general population using social media advertisements. Participants were required to be above 18 years of age, have no hearing impairments, and normal or corrected to normal vision, which was assessed by self-report. Participants were reimbursed with a gift card valued 200 SEK (approximately $25) on completion. The study was approved by the Swedish Ethical Review Authority and conducted in accordance with the Helsinki Declaration. All participants actively provided informed consent to participate in the study.

### Materials

Consent and questionnaire data were collected and managed using REDCap (Research Electronic Data Capture), a secure, web-based software platform designed to support data capture for research studies ^21,22^, hosted at Uppsala University. To assess trait anxiety, participants completed the Spielberger State-Trait Anxiety Inventory - Trait version (STAI-T) using the REDCap server ^23^. The fear conditioning experiment, Screaming Internet Lady, was delivered online through the PsyToolkit platform ^19,20^. For experimental stimuli, images of two female faces were selected from the FACES database ^24^. Neutral facial expressions were used as conditioned stimuli (CS) and fearful facial expressions in conjunction with a fearful scream served as the unconditioned stimulus (US). Reaction time probes were 440 Hz tones with a duration of 200 ms.

### Procedure

Participants were recruited through social media advertisements targeting adults (18 years and older) living in Sweden. A web link to a RedCap server with information about the study and registration was included in the advertisement. After actively providing informed consent to participate in the study, subjects answered screening questionnaires. Those passing inclusion (>18 years old) and exclusion (uncorrected vision or hearing impairments) criteria were provided a link to further questionnaires and the fear conditioning task. Participants completed all questionnaires and experiments through the internet.

Prior to commencing the fear conditioning task, participants were instructed to sit in an undisturbed place and wear headphones. They then individually set their computer sound volume to be unpleasantly loud (but not hurt their hearing) using a two-step procedure. First, they listened to a recording of a neutral text and set their sound-level to their preferred listening volume. After this, they listened to a test sound (an alarm-sound) and were instructed to set their computer volume to be markedly unpleasant but bearable when listening to this sound. They were then provided with the instructions to the fear conditioning task, that they were to see faces on the screen, might hear screams, and that they would occasionally hear a tone (the auditory probe) that signaled that they should press the spacebar as quickly as possible. The Screaming Internet Lady fear conditioning task is an online adaptation of the Screaming Lady task ^25^, consisting of three phases: acquisition, extinction, and reinstatement. The instruction to press the spacebar as quickly as possible when hearing the probe tone was repeated before the extinction and reinstatement phases in accordance with previous studies by Dirikx et al ^16,17^.

During fear acquisition, participants were presented with two neutral female faces serving as conditioned stimuli (CS), see Fig. 1. The CS were presented 16 times each in a sequential order. One of the cues (CS+) was presented for 4000 ms and always followed by an image of the same woman with a fearful facial expression and a scream for 1000 ms, serving as unconditioned stimulus (US). The scream was played at the intensity set by the participant as described above. The other cue (CS−) was presented for 5000 ms and never followed by the US. The two CSs were counterbalanced across participants. CS presentations were interspersed with fixation crosses in the center of the screen for a random amount of time (mean: 2500 ms, range: 2000-3000 ms, in steps of 100 ms). In 12 of the 16 of each CS presentation, a probe tone was played for 200 ms. Six of the probes for each CS were played 500 ms (early probe) and six 2500 ms (late probe) after CS onset and indicated to the participant to press the spacebar as quickly as possible. The omission of the probe in 25% of the trials and the two different timings of the probe were used to make their presentation unpredictable, see Fig. 1. Responses faster than 100 ms or slower than 1500 ms were discarded similar to previous studies using reaction time as an index of fear conditioning ^16,17^.

**Figure 1.**
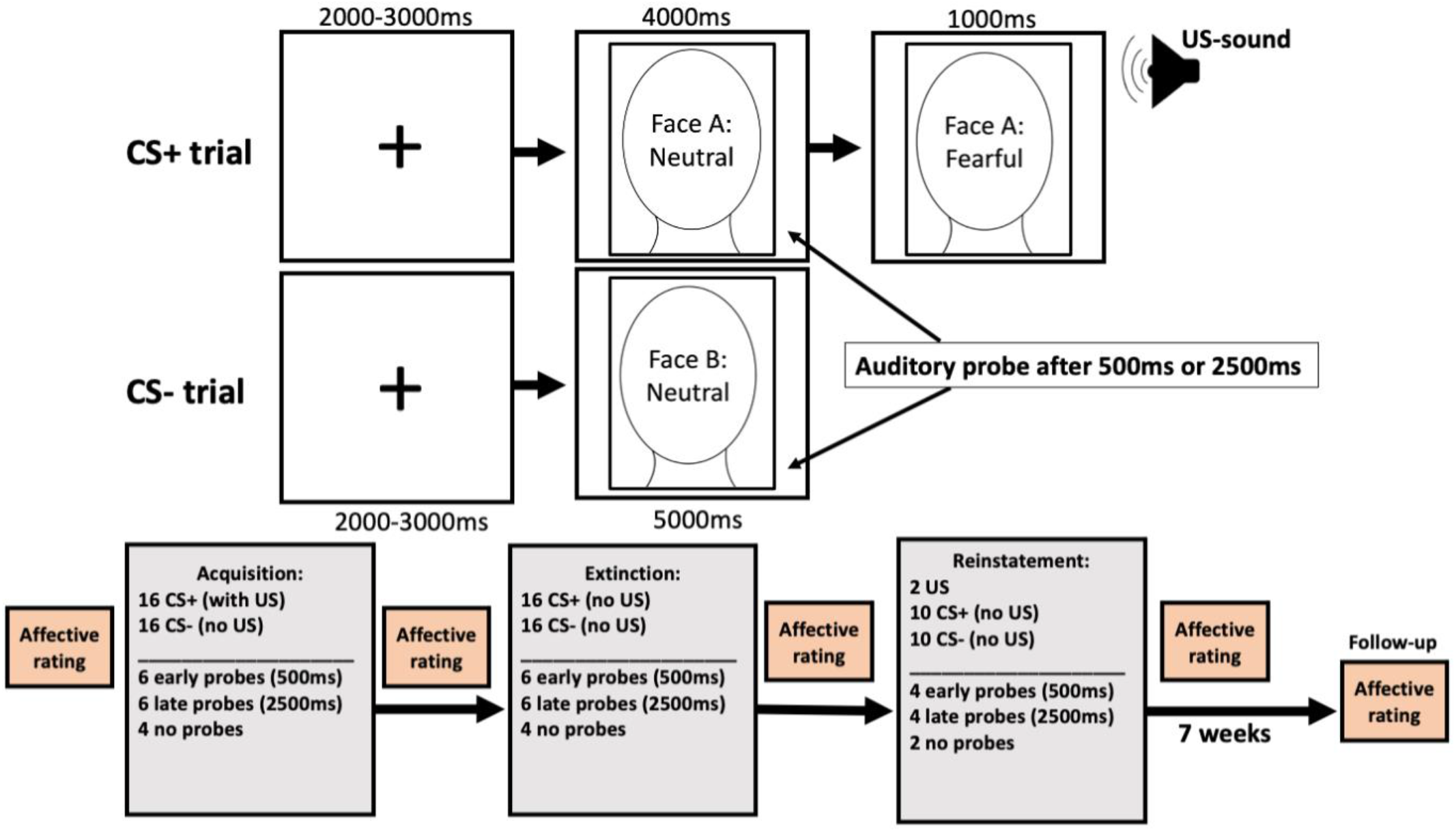
Experimental design. The top panel shows the outline of each experimental trial during the acquisition phase. A brief fixation cross was shown on the screen followed by an image of a neutral female face. During CS+ trials the presentation was followed by an aversive sound (scream) accompanied by the same face with a fearful expression. During CS- trials only the neutral expression was shown and no sound was played. During CS presentation, response times were probed after either 500ms or 2500ms by playing a brief 400Hz tone. Participants were instructed to press the space-bar as quickly as possible when the probe was played. Trials during extinction and reinstatement had the same structure but the US (sound and fearful expression) was always omitted. The face-images shown in the figure are not the same as the ones used in the experiment. The bottom panel illustrates the outline of the entire experiment, including number of trials of each type for each experimental phase.

The fear extinction phase was identical to the fear acquisition, but without the US. Thus, neither CS+ nor CS− were followed by the US during extinction. Reinstatement started with two unsignaled screams (but no fearful face) and then proceeded with 10 trials each of CS+ and CS−, in a sequential order, without presentation of the US. The probe tone was played in 8 of the trials for each CS (4 early and 4 late probes).

Before and after fear acquisition, after extinction and after reinstatement, participants rated how much they agreed from 1 (not at all) to 7 (totally) with feeling calm, fear, unpleasantness and irritation, while viewing the neutral CS faces. A composite rating score of the four feelings was created for each rating phase by reversing ratings on calm and then taking the mean of all ratings. After fear acquisition, participants rated how often (0 [never] - 100 [every time]) they heard a scream after viewing CS+ and CS−, used here as a measure of contingency awareness. After reinstatement, affective ratings to the US were collected using the same questions as for the CSs.

Participants were sent an e-mail approximately 4-5 weeks after the initial experiment, in which they were invited to complete the follow-up assessment. Reminder e-mails were sent out if they did not complete assessment. There was large variability in participants’ latency to complete the follow-up, and thus time from experiment to follow-up ranged from 5-15 weeks. Only affective ratings were collected at the follow-up.

### Missed responses and statistical analysis

In order to deal with missing responses for the response time measurements the following steps were taken. Subjects with incomplete data were excluded phase by phase according to the following principles: a maximum of 2 missed responses within each stimulus and probe-timing category, i.e. a maximum of 2 missed responses to early probes for the CS− during the acquisition phase, and so forth. As stated above, in addition to trials where the subject had not given a response, all trials with a response time below 100ms and above 1500ms were considered missed responses. In cases where participants had missed responses, but did not meet criteria for exclusion, imputation was performed according to the following principles: For each phase separately; in case of the first trial within a stimulus and probe-timing category was missing, the mean from the first trial of the other categories was imputed; in case of the last trial within a stimulus and probe-timing category was missing, the mean of the previous two trials was imputed; in other instances, the mean of the preceding and following trial within that stimulus and probe-timing category was imputed. For each of the phases none of the included subjects had more than 4 missed responses in total and the average number of missed responses per phase was 0.37, which we deemed acceptable. All statistical analyses were performed in JASP (version 0.14.1).

## Results

### US ratings and contingency awareness

As expected, participants rated the US high on negative affective ratings (M=5.7; SD=1.0; n=79, on the scale 1-7). Analysis of contingency ratings showed that participants were largely contingency aware. The average contingency rating for CS+ (M=67.4%; SD=32.4%; Median=75%; n=80) was substantially higher (t=12.96; df=79; p<.001) than CS− (M=8.6%; SD=19.6%; Median=0%; n=80), with 62 subjects (77.5%) rating at least 20 percentage points higher rate of screams following CS+ compared to the CS−.

### Affective ratings

To analyze the effect of fear conditioning on negative affective ratings, we entered rating values from the pre- and post-acquisition assessment points into a 2×2 repeated measures ANOVA with factors Phase (pre; acquisition) and Stimulus (CS+; CS−). A total of 80 subjects had complete data and were included in the analysis. The results revealed main effects of Phase (F=8.31; p=.005) and Stimulus (F=15.70; p>.001;) and crucially a Phase by Stimulus interaction effect (F=57.93; p<.001;) with a large effect-size (η^2^=0.110). Simple main effect analyses revealed a main effect of Stimulus after the acquisition phase (F=41.10; p<.001), where affective ratings of CS+ were higher compared to the CS−, but not during the pre-acquisition measurement (F=1.21; p=.275), where ratings were similar for the CS+ and CS−, see Fig. 2 and Table 1. Furthermore, simple main effects analyses showed main effects of Phase both for the CS+ category, where ratings increased from the pre to the post-acquisition measurement (F=42.24; p<.001), and also for the CS− where ratings decreased (F=4.66; p=.034), see Table 1. Thus, post-acquisition differences in negative affective ratings are not only driven by increases to the CS+ but also decreases to the CS−, possibly due to learning induced safety or relief associations, increasing positive valence of CS− stimuli. However, pre vs post acquisition effect-sizes were much larger for the CS+ (d=0.73) than for the CS− (d=−0.24).

**Table 1.**
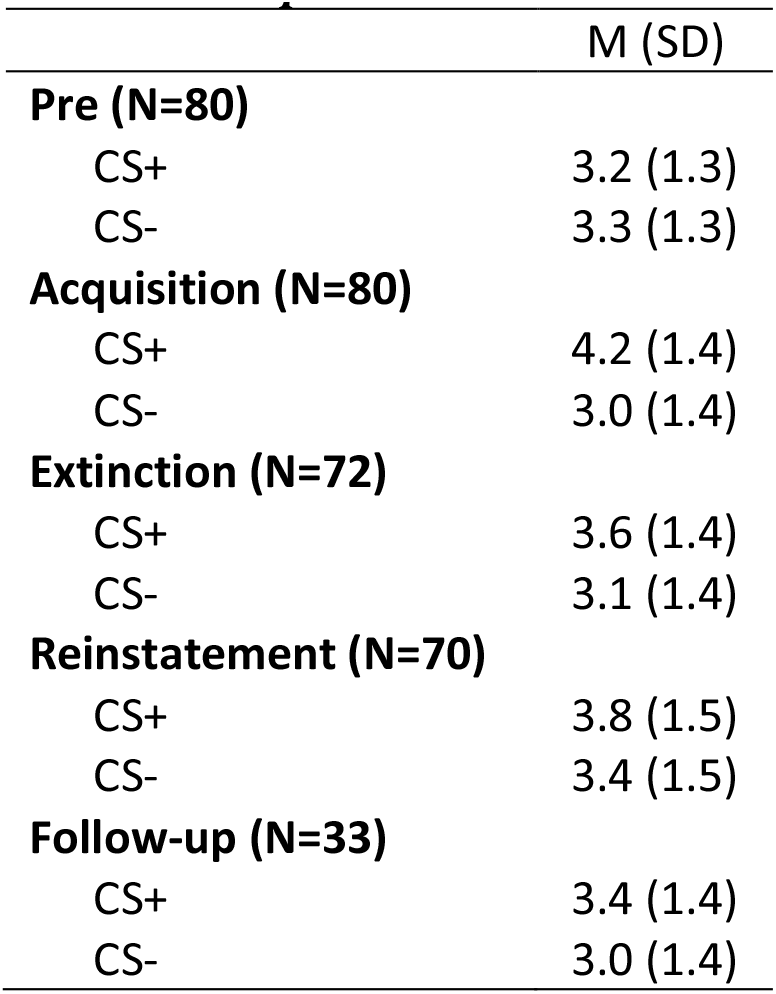
Descriptive statistics for negative affective ratings presented separately for each stimulus and phase.

|  | M (SD) |
| --- | --- |
| **Pre (N=80)** |  |
| CS+ | 3.2 (1.3) |
| CS- | 3.3 (1.3) |
| **Acquisition (N=80)** |  |
| CS+ | 4.2 (1.4) |
| CS- | 3.0 (1.4) |
| **Extinction (N=72)** |  |
| CS+ | 3.6 (1.4) |
| CS- | 3.1 (1.4) |
| **Reinstatement (N=70)** |  |
| CS+ | 3.8 (1.5) |
| CS- | 3.4 (1.5) |
| **Follow-up (N=33)** |  |
| CS+ | 3.4 (1.4) |
| CS- | 3.0 (1.4) |

**Figure 2.**
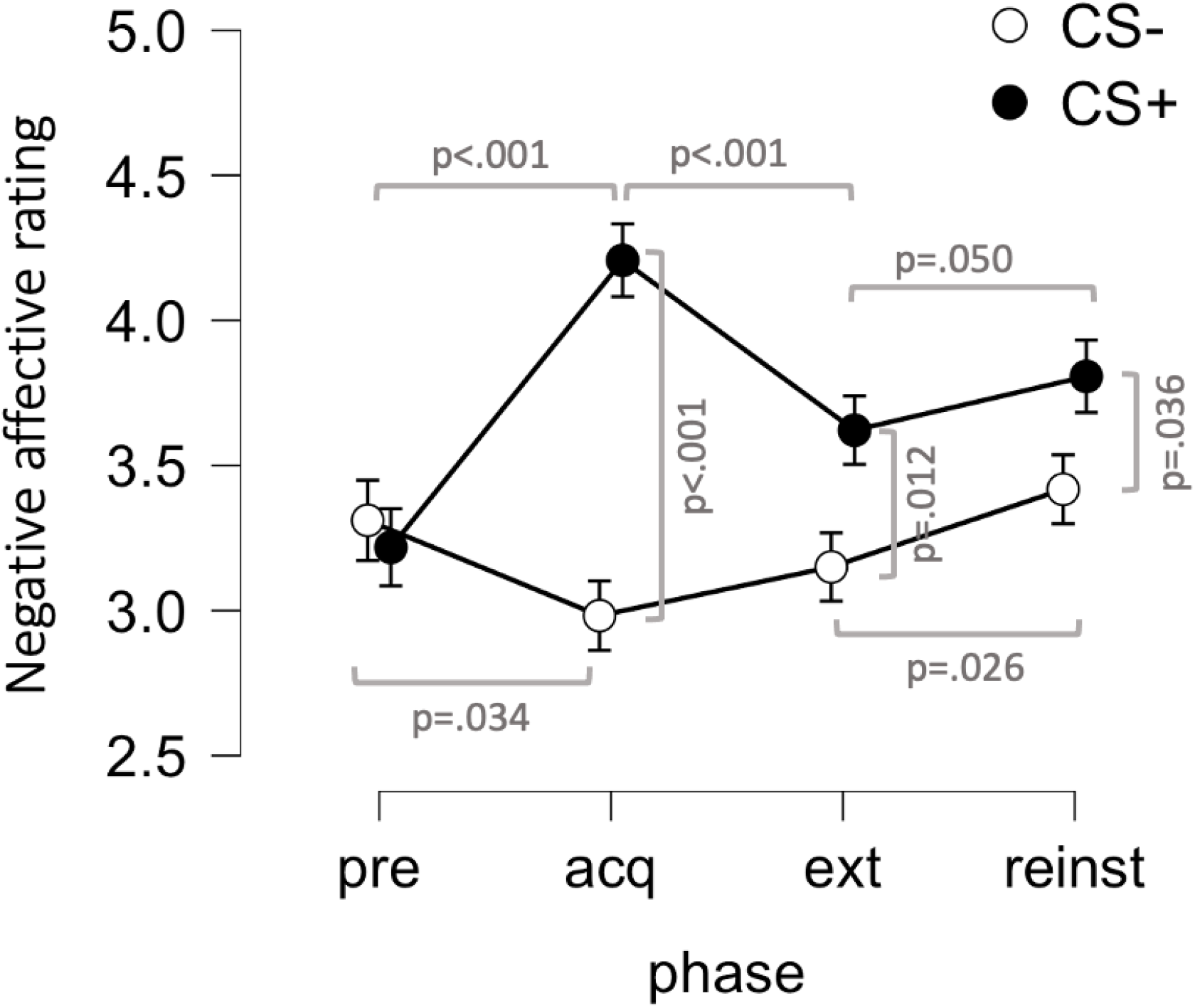
Affective ratings to conditioned stimuli (CS). Negative affective ratings were similar for stimuli paired with aversive outcome (CS+) and stimuli never paired with aversive outcome (CS−) pre acquisition, but differed after acquisition (acq), with increased negative ratings of CS+ and reduced negative ratings of CS−. Post extinction (ext), ratings of CS+ and CS− still differed, but to a lesser degree, whereas reinstatement (reinst) increased negative ratings of both CS. Affective ratings is a composite score consisting of the mean rating (1[not at all] to 7[totally]) of how much participants agree with feeling fear, discomfort, irritation and calm (reversed score). Points and error-bars indicate means and SEMs.

To analyze the effect of extinction on affective ratings, we entered rating values from the post-acquisition and post-extinction assessments into a 2×2 repeated measures ANOVA with factors Phase (acqusition; extinction) and Stimulus (CS+; CS−). Eight subjects were excluded from this analysis due to incomplete data during the extinction phase, thus 72 participants were included. The result showed main effects of Phase (F=7.24; p=.009) and Stimulus (F=24.97; p<.001) as well as a Phase by Stimulus interaction effect (F=23.44; p<.001). Simple main effects analyses showed a main effect of Stimulus both during post-acquisition (F=38.60; p<.001) as well as post-extinction (F=7.63; p=.007), although the effect was much larger after acquisition (d=0.72) compared to after extinction (d=0.33). Also, the main effect of Phase was only detected for the CS+ (F=30.04; p<.001) with significant decreases in negative affective ratings, whereas no effect of Phase was detected for the CS− (F=1.28; p=.261), where negative affective ratings were similar after acquisition and extinction, see Fig. 2 and Table 1. We found clear indications of extinction for affective ratings as indicated by the Phase by Stimulus interaction, and that this effect is driven by decreases to the CS+ whereas no change occurred for the CS−. Additionally, the current protocol does not appear to lead to complete extinction, since there was still an effect of Stimulus even after extinction.

To analyze possible effects of reinstatement, we entered rating values from the post-extinction and post-reinstatement assessment points into a 2×2 repeated measures ANOVA with factors Phase (extinction; reinstatement) and Stimulus (CS+; CS−). An additional 2 subjects were excluded from this analysis due to incomplete data after the reinstatement phase, thus 70 participants were included. The results showed main effects of Phase (F=7.54; p=.008) and Stimulus (F=6.47; p=.013), but no Stimulus by Phase interaction (F=0.38; p=.539). Simple main effects analyses showed main effects of Stimulus both after the extinction phase (F=6.73; p=.012) as well as the reinstatement phase (F=4.58; p=.036) with similar effect sizes (d=0.26 for CS+ vs CS− comparison after reinstatement). Also, there was a main effect of Phase, both for the CS+ (F=3.97; p=.050) and for the CS− (F=5.20; p=.026) with similar effect sizes (CS+: d=0.24; CS−: d=0.27 for Extinction vs Reinstatement comparisons), indicating significant increases in negative affective ratings for both stimuli. Thus, we found no clear evidence for reinstatement, as indicated by the absence of a Phase by Stimulus interaction, but rather that negative affective ratings tend to increase for both stimuli, and differences between CS+ and CS− are maintained.

Of the 70 participants who had complete data for acquisition, extinction and reinstatement, 33 (47%) completed the long-term follow-up assessment, which was performed approximately 5-10 weeks (M=52 days; SD=15 days; Min=35 days; Max=106 days) after the initial experiment. To investigate whether differential responding to experimental stimuli was still present, we adjusted the follow-up rating scores for baseline ratings (by the deducting the rating scores from the pre-assessment from the follow-up assessment for the CS+ and CS− respectively), and performed a t-test. The results showed a difference in rating-scores with a small to moderate effect-size (t=2.67; p=.012; df=32; d=0.46), where baseline-adjusted negative affective ratings were higher for the CS+ (M=0.5; SD=1.5) compared to the CS− (M=0; SD=1.5), indicating that the effect of learning on negative affective ratings was still present. Additionally, in order to evaluate whether ratings changed from the end of initial assessment point to the follow-up, we entered baseline-adjusted scores from the post-reinstatement phase and the follow-up phase into a 2×2 repeated measures ANOVA with factors Phase (Reinstatement; Follow-up) and Stimulus (CS+; CS−) only including participants with complete data (n=33). The results showed a main effect of Stimulus (F=5.37; p=.027) but no main effect of Phase (F=1.86; p=.182) and no Phase by Stimulus interaction (F=0.05; p=.821), indicating that neither negative affective ratings nor CS−differentiation changed during the 6-week follow-up period.

### Response times

To test whether response time could be used as a learning index for remote fear conditioning, we entered response times from the acquisition phase into a 2×2×6 repeated measures ANOVA with factors Stimulus (CS−; CS+), Probe-timing (early; late) and Trial (1-6). A total of 70 subjects had complete data for the acquisition phase and were included in these analyses. The results showed a main effect of Probe-timing (F=18.77; p<.001), where responses were slower for early probes (M=588ms; SD=157ms) compared to late probes (M=558ms; SD=156ms), a main effect of Trial (F=2.65; p=.033, Greenhouse-Geisser corrected), and a Stimulus by Probe-timing interaction (F=4.27; p=.043). No other effects could be detected (all Fs<2.11; all ps>.15). Analyses of simple main effects showed a main effect of Stimulus for early probes (F=4.84; p=.031) where responses to the CS+ where faster (M=575ms; SD=171ms) compared to the CS− (M=601ms; SD=157ms), but no main effect of Stimulus for late probes (F=0.002; p=.962) where response times were similar for the CS+ (M=558ms; SD=165ms) and the CS− (M=557; SD=158), see Fig. 2. Additionally, there was a clear main effect of Probe-timing for CS− (F=18.45; p<.001) but this effect was not robust for the CS+ (F=3.80; p=.055). This indicates that pairing a neutral stimulus with an aversive outcome shortens response time specifically to early probes, which could be used as a non-verbal learning index for remotely delivered fear conditioning protocols, although the effect size is small (d=0.26 for CS− vs CS+ during early probes).

For extinction, we focused on response times to early probes, since this was the only result that indicated learning effects during the acquisition phase. Thus, we compared average response times for the CS+ and CS− during the acquisition and extinction phases, restricted to early probes, using a 2×2 repeated measures ANOVA with factors Phase (acquisition; extinction) and Stimulus (CS−; CS+). Since 7 additional subjects had to be excluded due to incomplete data during the extinction phase, a total of 63 subjects were included in this analysis. The results showed a main effect of Phase (F=49.38; p<.001), where responses during acquisition (M=579ms; SD=152ms) where slower compared to extinction (M=491ms; SD=159ms), a main effect of Stimulus (F=4.72; p=.034), where responses across both phases were faster to the CS+ (M=526ms; SD=158ms) compared to the CS− (M=544ms; SD=144ms), but no Phase by Stimulus interaction (F=1.01; p=.319), see Fig 3. The main effect of Stimulus in combination with the absence of a Phase by Stimulus interaction effect indicates that extinction was not achieved by this protocol, however, simple main effects analyses showed a main effect of Stimulus only during acquisition (F=4.53; p=.037) and not during the extinction phase (F=0.54; p=.464). This suggests that the main effect of Stimulus is driven by differences during acquisition, and lends tentative support to an interpretation that the response time difference established during acquisition is reduced during extinction when the US is omitted.

**Figure 3.**
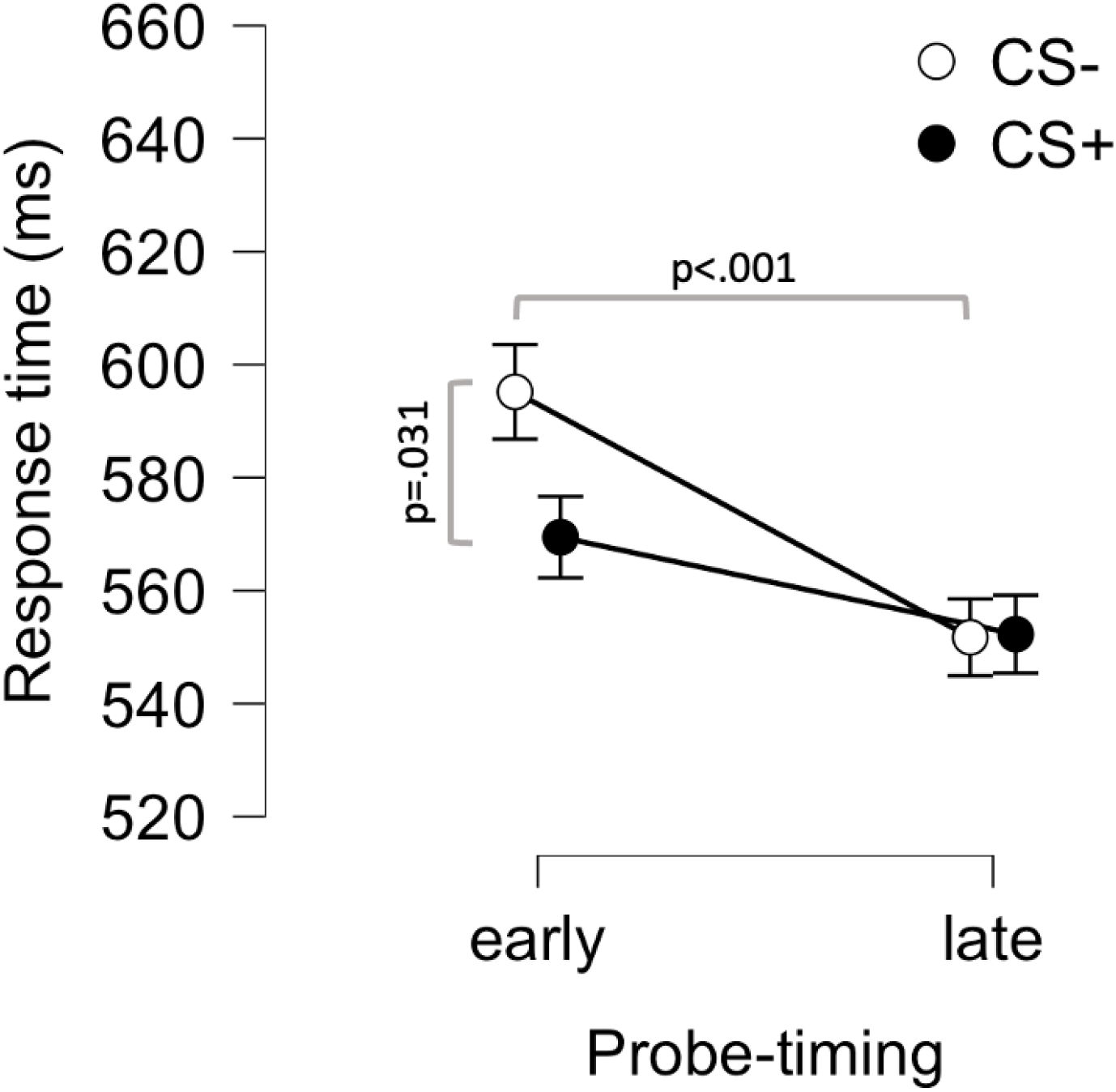
Response times to early and late auditory probes during presentation of conditioned stimuli (CS) in the acquisition phase. Response time to the neutral stimulus paired with an aversive outcome (CS+) was faster than to the stimulus never paired with an aversive outcome (CS−) during early (500 ms after onset of CS) but not late (2500 ms) probes. Points and error-bars indicate means and SEMs.

To evaluate possible effects of reinstatement we also focused on response times to early probes. We compared average response times for the CS+ and CS− during the extinction and reinstatement phases, to see if CS differentiation reemerges after presentations of 2 un-signaled US, using a 2×2 repeated measures ANOVA with factors Phase (extinction; reinstatement) and Stimulus (CS−; CS+). Two additional subjects had to be excluded due to incomplete data during the reinstatement phase, thus, 61 subjects were included in this analysis. The results did not show any main effects of either Phase (F=2.55; p=.116) nor Stimulus (F=0.11; p=.743), and no Phase by Stimulus interaction (F=0.36; p=.549). Consequently, we do not find any support that the protocol elicited reinstatement effects, but rather that response times during extinction and reinstatement are highly similar, and do not differ between the CS+ and CS−.

### Relations between fear conditioning and trait anxiety

#### Affective ratings

To explore possible associations with trait anxiety, as has been demonstrated previously using remotely delivered fear conditioning ^14^, we used the STAI-T to investigate correlations between this measurement and affective ratings of conditioned stimuli during the different phases. Thus, we calculated an acquisition index where ratings of each stimulus during the pre-assessment was subtracted from ratings after acquisition and also calculated a CS difference score (CS^diff^), where ratings of the CS− was subtracted from the CS+ adjusted for baseline-ratings. Similarly, we adjusted ratings of the CS+ and CS− post-extinction, by subtracting baseline ratings for each stimulus, and also calculated an extinction-index by subtracting CS^diff^ after extinction from CS^diff^ after acquisition. Pertaining to acquisition, the results showed a weak correlation between STAI-T and the baseline-adjusted CS− rating (r=0.22; CI^95%^=.004 to.423; p=.046; n=80), but we detected no correlation to the CS+ rating (r=0.11; CI^95%^= −.12 to.318; p=.351; n=80) or the CS^diff^ (r=−0.09; CI^95%^= −.30 to.13; p=.426; n=80), see Figure 4. For the extinction-phase we detected no correlations to the CS− (r=0.19; CI^95%^= −.05 to .40; p=.192; n=72), the CS+ (r=0.03; CI^95%^= −.20 to .26; p=.811; n=72), the CS^diff^ (r=−0.17; CI^95%^= −.38 to .07; p=.167; n=72) or the extinction-index (r=0.06; CI^95%^= −.18 to .29; p=.623; n=72). Although exploratory, we did find indications that specifically CS− ratings after acquisition is related to trait-anxiety, but this should be interpreted with caution.

**Figure 4.**
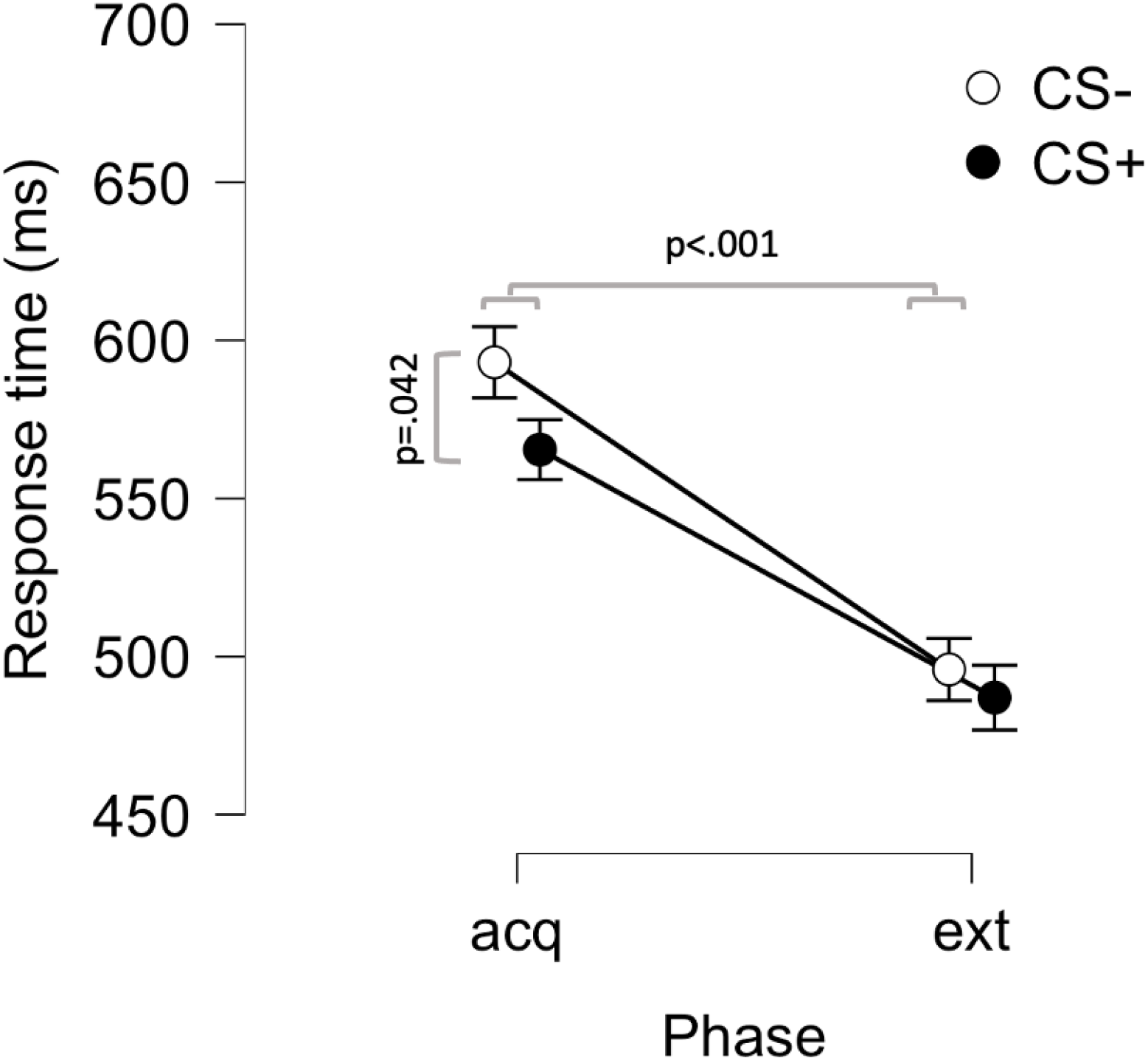
Response times to early auditory probes during presentation concomitant with conditioned stimuli (CS) in the acquisition (acq) and extinction (ext) phases. Response time to stimuli paired with an aversive outcome (CS+) was faster than to stimuli never paired with an aversive outcome (CS−) during acquisition but not extinction. Response times for both stimuli was higher during acuisition than during extinction Points and error-bars indicate means and SEMs.

**Figure 5.**
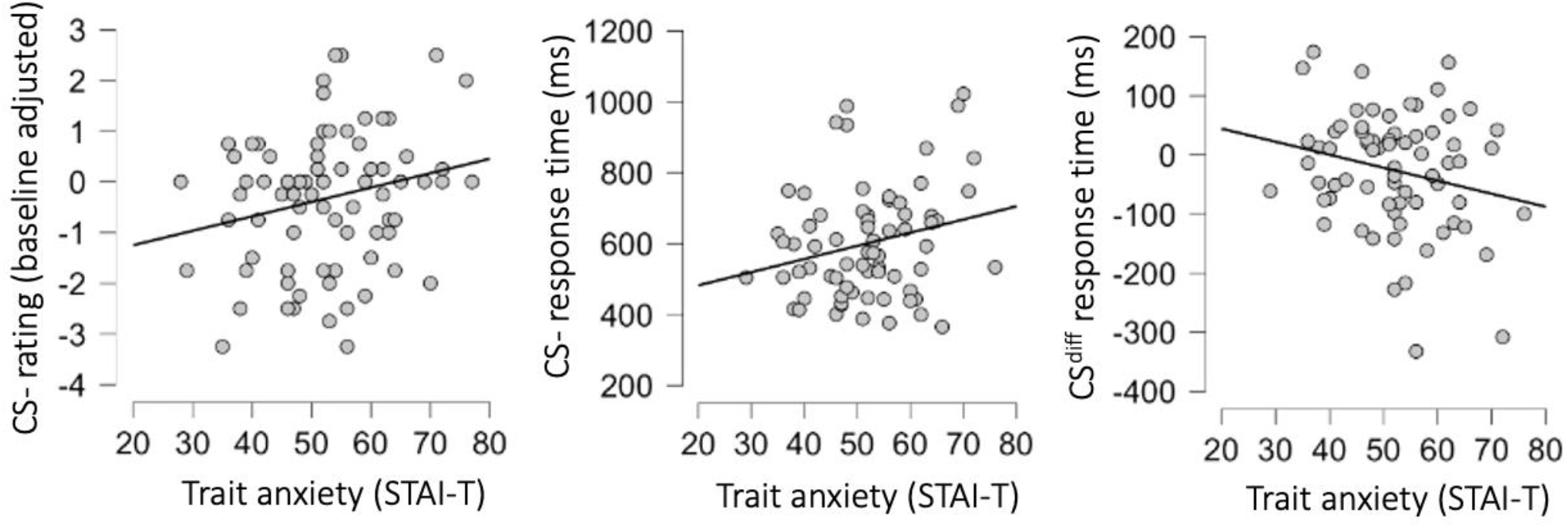
Associations between trait anxiety and learning variables. Scatter plots and linear fit lines showing correlations between trait anxiety, measured using Spielberger State-Trait Anxiety Inventory - trait version (STAI-T) and (left) negative affective ratings to conditioned stimulus never paired with an aversive outcome (CS−), (middle) response time to probes presented during CS−, and (right) difference in response time between probes presented during stimuli paired with and aversive outcome (CS+) and CS−. A negative difference score indicates that CS+ responses were faster than CS− responses, and thus reflects stronger learning.

#### Response times

We also performed similar analyses for the response time measure. Since we only found evidence of learning effects for early probes during acquisition, we restricted the analyses to this outcome. Thus, we investigated the relationship between STAI-T score and response time to the CS−, the CS+ and the CS^diff^-score for early probes only during acquisition. Similar to affective ratings we found a weak correlation between response time to the CS− and STAI-T (r=0.24; CI^95%^= .002 to .447; p=.048; n=70), no correlation to the CS+ (r=0.09; CI^95%^= −.15 to .32; p=.467; n=70) and an uncertain weak negative association to the CS^diff^-score score (r=−0.22; CI^95%^= −.43 to .02; p=.066; n=70), see Figure 4. Note, that given the direction of the effect of learning on response times, where CS+ responses are faster than CS− responses, a negative CS^diff^-score indicates stronger learning. Thus, the negative correlation for CS^diff^ suggests that stronger learning is related to higher scores on the trait-anxiety measure.

## Discussion

In this internet-based study of fear conditioning, we first replicate and extend previous findings of increased negative affective ratings to a cue (CS+) predicting an aversive scream. Next, we also demonstrate that negative affective ratings to CS− (predicting no aversive scream) are reduced during fear acquisition. Finally, we found tentative support for stimulus concurrent reaction time as an index of fear learning. However, the latter findings were weak and should be replicated before drawing firm conclusions about the feasibility of this approach to index cued fear responses non-verbally during remotely delivered fear conditioning.

Concerning negative affective ratings, essentially, our results replicate a previous findings on remote fear conditioning ^14^, in that we see clear indications of learning using affective ratings of conditioned stimuli. We also extend previous findings, showing that differential responding to conditioned stimuli post-acquisition is driven by both increases to CS+, reflecting fear learning, as well as decreases to CS−, possibly reflecting safety learning or relief conditioning. We also find strong evidence for extinction using affective ratings, which could not be established by Purves et al. ^14^, since they only recorded affective ratings pre acquisition and post extinction. The current results demonstrate the feasibility of investigating questions related to extinction processes using remote fear conditioning and affective ratings. We did not, however, find evidence of reinstatement using the current methodology, but rather a general increase in negative affective ratings after unsignaled US presentations. It should be noted that our study was poorly designed to detect such effects because affective ratings were collected only after the whole reinstatement phase, which included 2 un-signaled US presentations and 8 trials of each CS. Thus, the additional non-reinforced trials may have caused further extinction and attenuated any effect of US presentation on affective ratings. This could have been mitigated by using trial-by-trial affective ratings or rating sooner after the US presentations ^9^. Furthermore, when investigating reinstatement, a between-subject design would have been much preferred in order to isolate the effects of un-signaled US presentation. Pertaining to results from the follow-up test, we could show that effects on affective ratings was still present, even after six weeks or more. This supports the conclusion that the experimental protocol elicits long-term retention. We could not, however, find any support for spontaneous recovery since the affective ratings and CS−difference scores were largely unchanged from the end of the initial experimental session to the follow-up.

We could not replicate Purves et al. ^14^ findings of an association between CS+ affective ratings post-extinction and trait anxiety, but instead identified such an association for post-acquisition affective ratings of CS−, in that higher trait anxiety was associated with higher ratings of negative affect for the CS− and not the CS+. Our findings are exploratory and should be interpreted with caution, but they are in line with previous lab-based studies that have found similar results using trial by trial distress-ratings during fear conditioning ^9^, and also with studies on clinical populations that have shown enhanced fear responses to safety stimuli in patients with anxiety disorders using various outcome measurements ^6^. Future studies employing remote fear conditioning, with larger sample sizes could possibly lead to more firm conclusions regarding this effect.

The second aim of our study was to test the feasibility of using concurrent measures of simple reaction time to an auditory probe during CS presentation as a learning-index for aversive Pavlovian conditioning and extinction. To this regard, we only found weak evidence that the current fear conditioning protocol has an impact on response time. Effects were small and could only be observed for probes presented early during CS presentation and only in the acquisition phase. Furthermore, effects were opposite to the predicted direction, as previous lab-based studies investigating this effect ^16,17^ observed slower response times to CS+ using a similar protocol in a lab-environment, and interpreted this result as an effect of attention capture by the visual CS+. We instead found faster response time to early probes during CS+ presentation, which would be more in line with an interpretation that exposure to an aversively conditioned stimulus leads to a rapid but quickly dissipating increase in vigilance. This could be construed as evolutionary adaptive, since it would allow the organism to respond faster to possible environmental threats. Note that previous lab-based studies using other experimental set-ups have found similar effects of faster response times to the CS+ compared to the CS− ^26–29^ but results have been inconsistent with other studies showing no effects or effects in the opposite direction ^13^. In addition, we only found tentative support that conditioned response time modulation undergoes extinction. Overall, we view our results as inconclusive with regard to the question of whether response times can be used as a non-verbal learning index for remote fear conditioning and extinction experiments. We did find differences in response time to conditioned stimuli during acquisition, but the effects were weak, suggesting that protocols that lead to more robust effects need to be developed in order for this experimental paradigm to be useful.

Similar to our findings of affective ratings, we also found an association between response times to the CS− and trait anxiety, as well as an uncertain association between response time CS^diff^-score and trait anxiety. Although an association between responses to the safety cue and trait-anxiety could be expected based on previous studies ^6,9^, the direction of this effect is surprising. We found that slower responses to the CS− was associated with higher trait-anxiety also an uncertain negative association to the CS^diff^-score, but since the conditioning effect indicated that responses are faster to the CS+, this means that stronger fear learning (as measured here) is associated more trait anxiety. This runs counter to previous studies showing weaker fear learning in pathological anxiety due to enhanced responding to safety cues ^6^. This finding is exploratory and should be interpreted with caution. Furthermore, much larger sample-sizes are likely required in order to reliably detect an association of the current magnitude. If replicated however, this could prove to be an interesting finding in that it would show that conditioned response time modulation is differently associated to pathological anxiety compared to other CR-estimation methods, which could shed light on how fear learning is related to problematic anxiety.

Some limitations of our study should be mentioned. First, there is a large majority of women in the sample, which may hamper generalization of the findings to men. Second, participants performed the fear conditioning task online in their own homes or in other locations where we did not have control over their environment or compliance. Nonetheless, participants were instructed to complete the task in an undisturbed location and they indicated in questions after the conditioning paradigm that they had performed the paradigm in accordance with instructions. Furthermore, although we only found weak evidence for the response time outcome, we found strong effects for affective ratings in line with previous findings, which indicates that compliance was adequate to detect conditioning effects.

In conclusion, the internet-based fear conditioning paradigm described in this study is suitable for investigating remotely delivered fear acquisition and extinction using affective ratings as outcome in large samples in a cost-effective manner. More research is needed to find optimal parameters for probe presentation and experimental design before reaction time can be reliably employed in large-scale online studies of fear conditioning. The current study constitutes a first step in this direction.

## Acknowledgements

We thank all participants without whom we could not have performed this research. The study was supported by funding from Kjell och Märta Beijers stiftelse, Magnus Bergvall Foundation, The Swedish Brain Foundation, Riksbankens Jubileumsfond, The Swedish Research Council and the Craaford foundation. Dr. Pine is supported by National Institute of Mental Health (NIMH) Intramural Research Program (IRP) Project ZIA-MH002781.

